# Flexible cobamide metabolism in *Clostridioides (Clostridium) difficile* 630 Δ*erm*

**DOI:** 10.1101/772582

**Authors:** Amanda N. Shelton, Xun Lyu, Michiko E. Taga

## Abstract

*Clostridioides (Clostridium) difficile* is an opportunistic pathogen known for its ability to colonize the human gut under conditions of dysbiosis. Several aspects of its carbon and amino acid metabolism have been investigated, but its cobamide (vitamin B_12_ and related cofactors) metabolism remains largely unexplored. *C. difficile* has seven predicted cobamide-dependent metabolisms encoded in its genome in addition to a nearly complete cobamide biosynthesis pathway and a cobamide uptake system. To address the importance of cobamides to *C. difficile*, we studied *C. difficile* 630 Δ*erm* and mutant derivatives under cobamide-dependent conditions *in vitro*. Our results show that *C. difficile* can use a surprisingly diverse array of cobamides for methionine and deoxyribonucleotide synthesis, and can use alternative metabolites or enzymes, respectively, to bypass these cobamide-dependent processes. *C. difficile* 630 Δ*erm* produces the cobamide pseudocobalamin when provided the early precursor 5-aminolevulinc acid or the late intermediate cobinamide, and produces other cobamides if provided an alternative lower ligand. The ability of *C. difficile* 630 Δ*erm* to take up cobamides and Cbi at micromolar or lower concentrations requires the transporter BtuFCD. Genomic analysis revealed genetic variations in in the *btuFCD* locus of different *C. difficile* strains, which may result in differences in the ability to take up cobamides and Cbi. These results together demonstrate that, like other aspects of its physiology, cobamide metabolism in *C. difficile* is versatile.

**Importance:** The ability of the opportunistic pathogen *Clostridioides difficile* to cause disease is closely linked to its propensity to adapt to conditions created by dysbiosis of the human gut microbiota. The cobamide (vitamin B_12_) metabolism of *C. difficile* has been underexplored, though it has seven metabolic pathways that are predicted to require cobamide-dependent enzymes. Here, we show that *C. difficile* cobamide metabolism is versatile, as it can use a surprisingly wide variety of cobamides and has alternative functions that can bypass some of its cobamide requirements. Furthermore, *C. difficile* does not synthesize cobamides *de novo*, but produces them when given cobamide precursors. Better understanding of *C. difficile* cobamide metabolism may lead to new strategies to treat and prevent *C. difficile-*associated disease.

## Introduction

The human gut microbiota is a complex community composed of hundreds to thousands of species of bacteria, archaea, and eukaryotic microbes (1). Members of this community compete for nutrients such as carbon sources, but also release metabolites that benefit other members. The exchange of B vitamins, particularly vitamin B_12_, is thought to be prevalent in many environments because most bacteria lack the ability to synthesize some of the cofactors they require for enzyme catalysis (2–6), and instead must acquire them from other organisms (7). Such nutrient cross-feeding interactions can influence bacterial metabolism in ways that can affect not only the microbiota, but also host health (8, 9).

*Clostridioides (Clostridium) difficile* is a human intestinal pathogen that is among the most common causes of nosocomial infections, with nearly 300,000 healthcare-associated cases per year in the United States (10). *C. difficile* colonization of the gut is correlated with dysbiosis of the gut microbiota (11). Its abilities to germinate from spores, proliferate in the gut, and cause disease are impacted both positively and negatively by ecological and metabolic factors (12–14). The global alteration of the gut metabolome following antibiotic treatment is correlated with increased susceptibility to *C. difficile* infection, and recent work has linked changes in the relative abundance of specific metabolites to changes in the microbiome using model systems (11, 15–17). For example, succinate availability increases after disturbance of the microbiota, allowing *C. difficile* expansion in a mouse model (18). Additionally, specific commensal bacteria have been shown to produce compounds that stimulate *C. difficile* metabolism. In a biassociation, *Bacteroides thetaiotaomicron* can break down host mucin and produce sialic acid, which can be used by *C. difficile* for expansion in the gut (19). *C. difficile* can also induce other members of the microbiota to produce indole, which is thought to create a more favorable environment for the pathogen by inhibiting competing microbes (20).

Some interactions with microbiota members have also been shown to be inhibitory to *C. difficile*. Co-culturing with certain *Bifidobacterium* spp. on particular carbon sources reduces *C. difficile* toxin production relative to monoculture (21). While primary bile acids produced by the host promote *C. difficile* spore germination, *Clostridium scindens* and other 7α-dehydroxylating Clostridia transform these compounds into secondary bile acids, which are inhibitory to *C. difficile* (22, 23). The latter example illustrates that compounds in the same class can have different effects on the disease state. Given the complexity of metabolic interactions in the mammalian gut, many additional microbial metabolites likely influence the ability of *C. difficile* to colonize and persist in the gut.

One class of metabolites that has not been explored for its ability to affect *C. difficile* growth and virulence is cobamides, the vitamin B_12_ (also called cobalamin) family of cofactors. Cobamides are used in diverse microbial metabolisms including methionine synthesis, deoxyribonucleotide synthesis, acetogenesis, and some carbon catabolism pathways. These reactions are facilitated by fission of the Co-C bond to the cobamide upper ligand, which can be a 5’-deoxyadenosyl group for radical reactions, a methyl group for methyltransferase reactions, or a cyano group in the inactive vitamin form (24) (labeled as “R” in Fig. 1A). Over 80% of all sequenced bacteria (25–27) and 80% of sequenced human gut bacteria (2, 28, 29) have one or more cobamide-dependent enzymes, suggesting that cobamides are widely used cofactors across microbial ecosystems. Strikingly, fewer than 40% of bacterial species are predicted to produce cobamides *de novo* (2, 25–28), and therefore over half of bacteria that use cobamides must acquire them from their environment. Cobamides vary in the structure of the lower ligand (Fig. 1A, B), and organisms studied to date are selective in which cobamides they can use (28, 30–37). Seven cobamides in addition to the cobamide precursor cobinamide (Cbi, Fig. 1C) have been detected in the human gut (38). In an environment with plentiful, diverse cobamides and cobamide precursors, a microbial species that requires a particular cobamide can either import that cobamide, synthesize it *de novo*, chemically remodel available cobamides to the preferred structure, or alter its need for the cobamide by using alternative pathways (8, 39).

**Figure 1.**
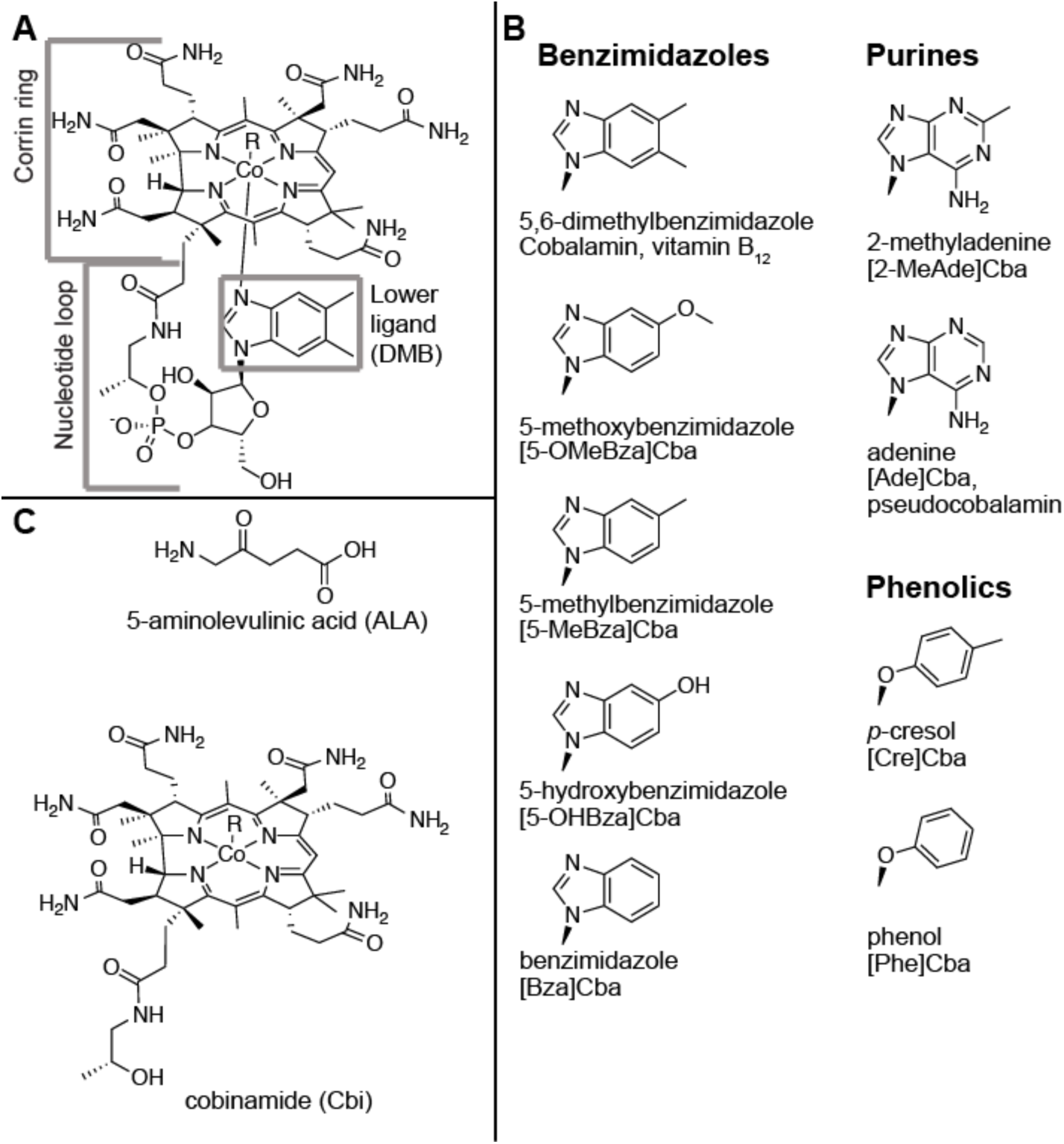
Structures of cobamides and cobamide precursors. **A.** Structure of cobalamin (B_12_). The corrin ring, nucleotide loop, and lower ligand are labeled. **B.** Lower ligands of cobamides analyzed in this study, with the three structural classes labeled. The lower ligand name, abbreviation for the cobamide containing the lower ligand, and alternative names of the cobamide (when applicable) are indicated. **C**. Cobamide precursors used in this study. R, upper ligand (-CN, -OH, -CH_3_ or 5’-deoxyadenosyl).

The seven predicted cobamide-dependent enzymes encoded in the *C. difficile* genome are involved in methionine synthesis, nucleotide metabolism, and carbon metabolism (Fig. 2). When grown with amino acids and glucose as carbon and energy sources *in vitro, C. difficile* does not require cobalamin supplementation (40). However, in model infection systems, cobamide-dependent metabolism may be important for virulence and growth. For example, access to ethanolamine catabolism may be important in modulating virulence, as deletion of EutA, the reactivating factor required for activity of the cobamide-dependent ethanolamine ammonia lyase (EutBC), in *C. difficile* strain 630 Δ*erm* reduces the mean time to morbidity in a hamster model (41). Additionally, metabolic models and transcriptomics (42, 43) suggest that the cobamide-dependent Wood-Ljungdahl carbon fixation pathway is an important electron sink, and an experimental study suggests that it may be used for autotrophic growth by some *C. difficile* strains (44).

**Figure 2.**
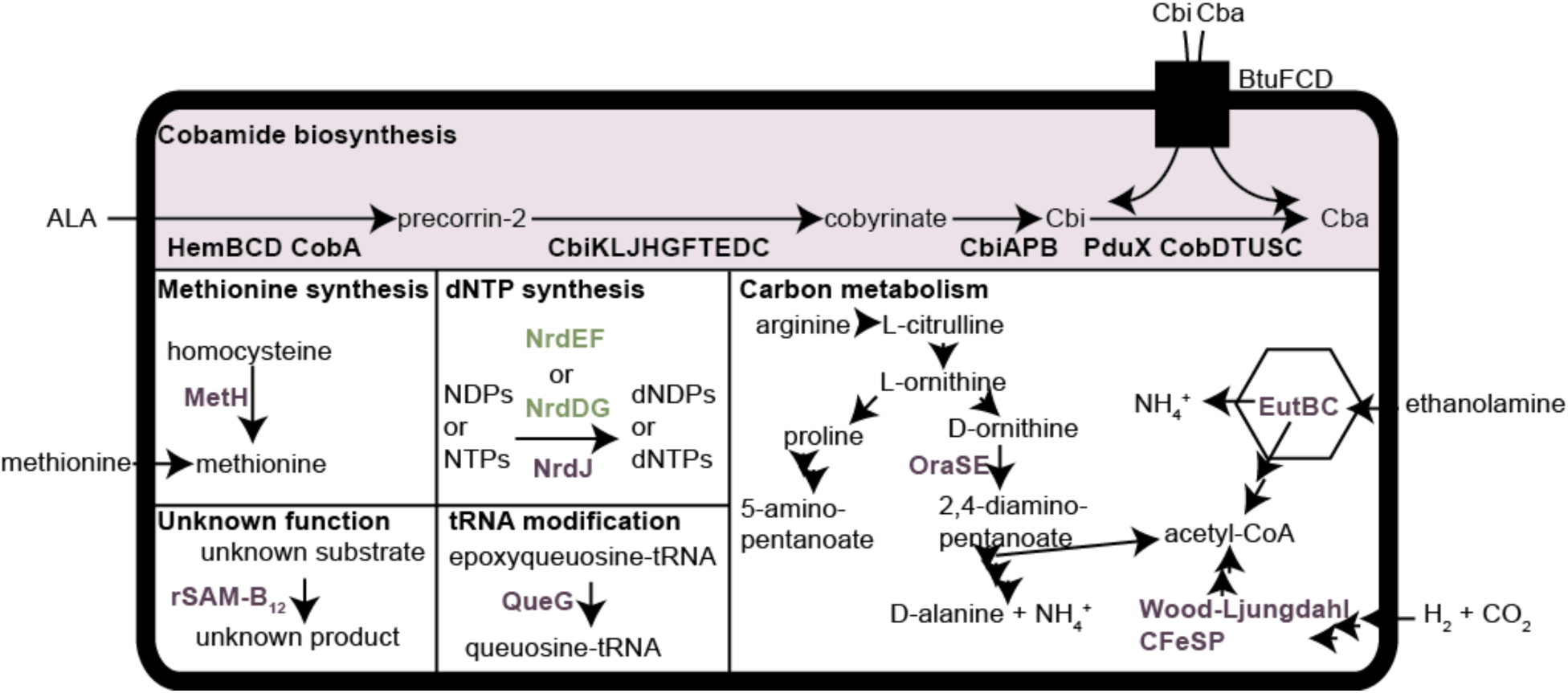
Predicted cobamide metabolism in *C. difficile* 630 Δ*erm*. The cobamide biosynthesis pathway is shown with a purple background, homologs of cobalamin-dependent enzymes in purple text, cobalamin-independent isozymes in green text, and the transporter BtuFCD as a black rectangle. Abbreviations: Cba, cobamide; Cbi, cobinamide; ALA, 5-aminolevulinic acid; rSAM, radical *S*-adenosylmethionine; NDPs, ribonucleoside diphosphates; NTPs, ribonucleoside triphosphates; dNDPs, deoxyribonucleoside diphosphates; dNTPs, deoxyribonucleoside triphosphates. Enzymes: MetH, cobalamin-dependent methionine synthase; NrdEF, cobalamin-independent, aerobic (oxygen-requiring, class I) ribonucleotide reductase (RNR); NrdDG, cobalamin-independent, anaerobic (oxygen-sensitive, class III) RNR; NrdJ, cobalamin-dependent (class II) RNR; QueG, epoxyqueuosine reductase; EutBC, ethanolamine ammonia lyase; CFeSP, corrinoid iron-sulfur protein; OraSE, D-ornithine 4,5-aminomutase.

The observation that *C. difficile* can grow without added cobamides *in vitro* (40) suggests that it may not require cobamides under those conditions, or that it can biosynthesize cobamides. However, all sequenced strains of *C. difficile* are missing *hemA* and *hemL*, the first two enzymes in the cobamide biosynthesis pathway required for production of the precursor 5-aminolevulinic acid (ALA) (45) (Fig. 1C). Therefore, *C. difficile* is predicted to be able to produce a cobamide only when ALA is available, as has been observed in three other bacteria (25) (Fig. 2). In order to use cobamide-dependent metabolisms, we predict that *C. difficile* requires cobamides or precursors such as ALA from the gut. While ALA is an intermediate made in all tetrapyrrole-producing organisms, including the host, cobamides are only produced by some bacteria and archaea (46).

To address the importance of cobamides for *C. difficile* metabolism and to understand how *C. difficile* acquires cobamides, we examined *C. difficile* 630 Δ*erm* and mutant derivatives *in vitro* under cobamide-dependent conditions. We found that the bacterium can use a surprisingly diverse array of cobamides for methionine and deoxyribonucleotide synthesis, and can use alternative nutrient sources or enzymes to fulfill its metabolic needs. In addition to importing and using a variety of cobamides, when provided with ALA or the late intermediate Cbi, *C. difficile* 630 Δ*erm* can produce the cobamide pseudocobalamin, and can produce other cobamides if provided an alternative lower ligand. Together, these results show that *C. difficile* is versatile in its cobamide metabolism.

## Results

### *C. difficile* requires methionine or a cobamide for growth

To investigate cobamide-dependent metabolism in the model *C. difficile* strain 630 Δ*erm*, we sought to culture the organism in conditions that require specific cobamide-dependent enzymes. The *C. difficile* genome encodes the cobalamin-dependent methionine synthase MetH, but does not contain the cobalamin-independent alternative enzyme MetE. The absence of a complete cobamide biosynthesis pathway suggests that *C. difficile* requires either methionine or a cobamide in its growth medium. Previously, methionine was classified as a “growth-enhancing,” but not essential, amino acid in a medium containing cyanocobalamin (vitamin B_12_) for seven of eight strains tested (40, 47). To test whether *C. difficile* can use cobamides for methionine synthesis and to identify the specific cobamides that support its MetH-dependent growth, we cultured *C. difficile* in a defined medium lacking methionine with a range of concentrations of cyanocobalamin, Cbi, and eight other cyanylated cobamides that we purified. *C. difficile* was unable to grow in this medium without cobamide or methionine addition (Fig. 3A), suggesting that, as predicted, it cannot produce cobamides *de novo* to support the activity of MetH. Remarkably, unlike other bacteria that have been reported to use a limited number of cobamides for methionine synthase activity (28, 48, 49), all of the cobamides and Cbi were able to confer high growth yields to *C. difficile* at concentrations as low as 1 nM (Fig. 3A). Methionine addition also supported growth, though higher concentrations were required than for cobamides (Fig. 3B). We also observed robust growth with addition of ALA (Fig. 3C).

**Figure 3.**
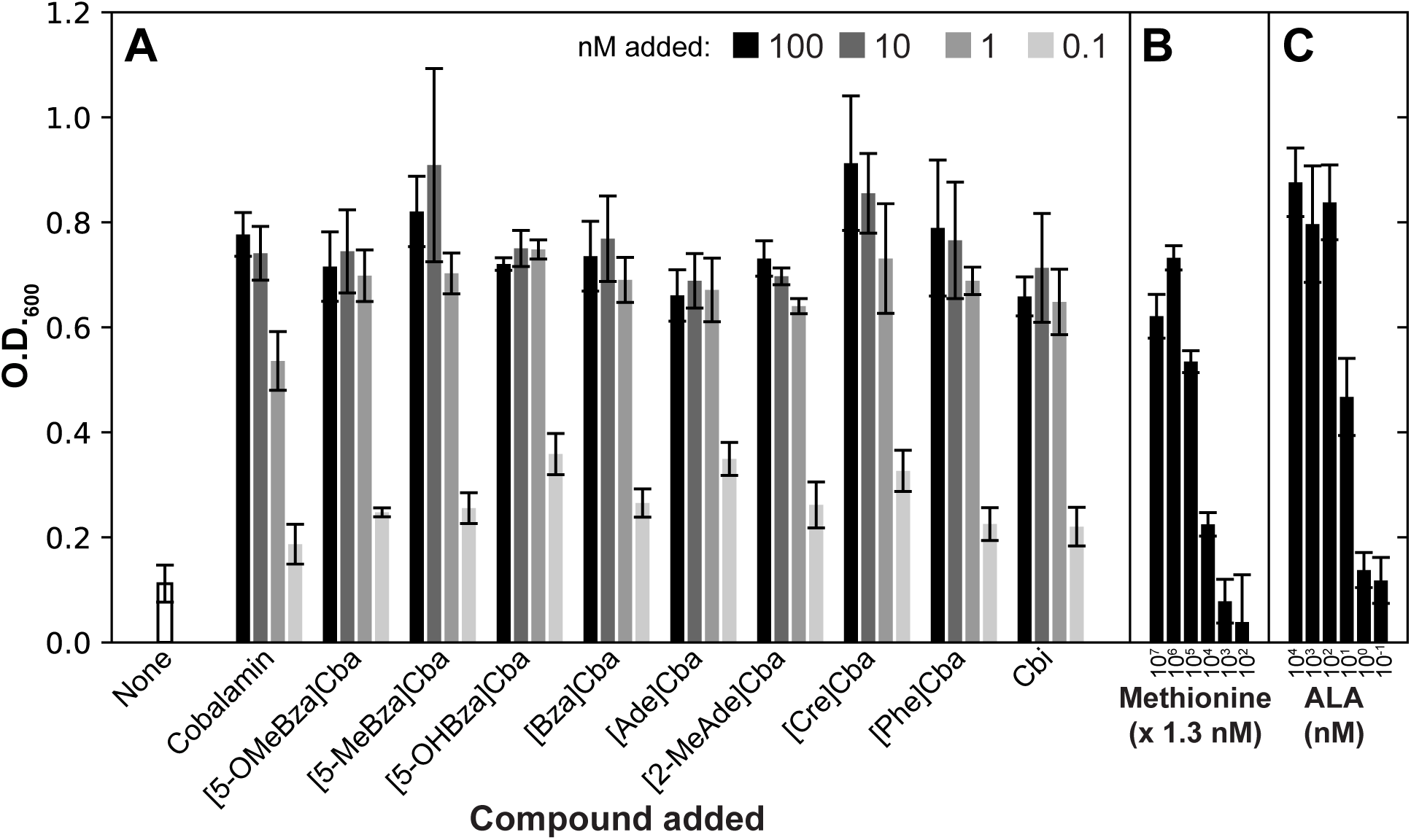
*C. difficile* can use a broad range of cobamides for MetH-dependent growth. The O.D._600_ of *C. difficile* 630 Δ*erm* cultures grown to saturation (22.5 hours) in CDDMK medium plus glucose without methionine with the addition of **A.** cobamides or Cbi, **B.** methionine, and **C.** ALA is shown. The mean and standard deviation of four biological replicates are shown in the bars and error bars, respectively.

### *C. difficile* growth with ribonucleotide reductase NrdJ requires a more restricted set of cobamides

*C. difficile* genomes encode homologs of the cobalamin-dependent (class II) ribonucleotide reductase (RNR) (*nrdJ*, CDIF630erm_RS07280), as well as two cobalamin-independent RNRs: an oxygen-dependent (class I) RNR (encoded by *nrdE*, CDIF630erm_RS16325, and *nrdF*, CDIF630erm_RS16320) and an oxygen-sensitive (class III) RNR (*nrdD*, CDIF630erm_RS00990 and *nrdG*, CDIF630erm_RS00995). In principle, any of these three isozymes could be used for deoxyribonucleotide synthesis from ribonucleotides, although under anaerobic conditions only the class II and class III RNRs are expected to function. Cobamide addition is not required for anaerobic growth of the parent strain *C. difficile* 630 Δ*erm* Δ*pyrE* in a casamino acid medium (CDDM) with glucose, and adding cobamides or cobamide precursors did not affect growth yield (Supplemental Fig. 1), suggesting that the class III RNR, NrdDG, is functional under these conditions. To test whether the class II RNR, NrdJ, is functional, we deleted the *nrdD* and *nrdG* genes while providing exogenous cobalamin, using the allelic exchange system in a Δ*pyrE* background (50). This strain could grow only with cobalamin addition, suggesting that NrdJ is functional and NrdEF is not under these growth conditions (Fig. 4A). To determine which cobamides it requires, the Δ*nrdDG* strain was grown with the same cobamides and precursors as in Fig. 3. In contrast to MetH, NrdJ is more selective in the cobamides it can use (Fig. 4A), as expected based on studies with other class II RNRs (33, 36, 51, 52). There was little growth with [Cre]Cba, [Phe]Cba, and [5-OHBza]Cba (Fig. 4A). Addition of ALA also supported NrdJ-dependent growth (Fig. 4B).

**Figure 4.**
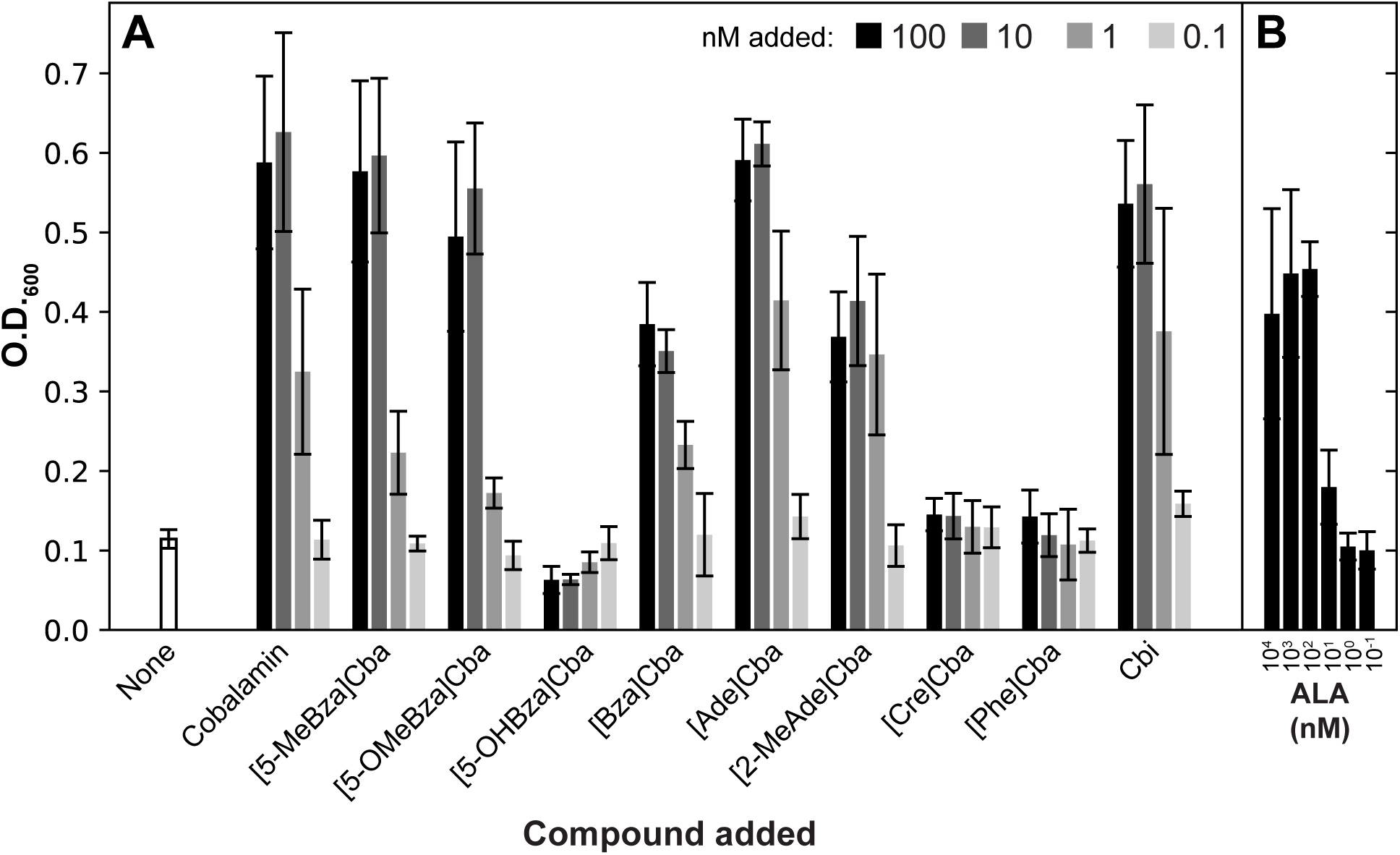
*C. difficile* is selective in which cobamides it can use for NrdJ-dependent growth. The O.D._600_ of *C. difficile* 630 Δ*erm* Δ*pyrE* Δ*nrdDG* cultures grown to saturation (22.5 hours) in CDDM with added uracil and glucose is shown for **A.** cobamides and Cbi, and **B.** ALA added. The mean and standard deviation of three biological replicates are shown in the bars and error bars, respectively.

### *C. difficile* produces pseudocobalamin from the precursor ALA via the *cbi* genes

The observation that *C. difficile* could grow under cobamide-dependent conditions with ALA or Cbi (Fig. 3A, C, Fig. 4) suggests that it can produce a cobamide from these precursors using the cobamide biosynthetic genes encoded in its genome (25). To test this prediction, the corrinoid fraction, which includes cobamides and late cobamide precursors including Cbi, was extracted from the cell pellets of *C. difficile* 630 Δ*erm* grown with either ALA or Cbi. Consistent with our predictions, HPLC analysis of the extracted corrinoids showed that *C. difficile* produced a cobamide only when ALA or Cbi was added (Fig. 5A). Additionally, as predicted, corrinoid analysis of a strain lacking the corrin ring biosynthesis genes *cbiKLJHGFTEDC* demonstrated that these genes are necessary for cobamide synthesis from ALA, but not Cbi (Fig. 5A). Because *C. difficile* lacks all known genes for biosynthesis of benzimidazoles and attachment of phenolic lower ligands, it is predicted to be incapable of producing benzimidazolyl or phenolyl cobamides, but may produce a purinyl cobamide (49, 53–59). Indeed, the major cobamide present in *C. difficile* corrinoid extracts co-eluted with the purinyl cobamide pseudocobalamin (Fig. 5A). The UV-Vis spectrum of the major cobamide was consistent with a pseudocobalamin standard (Supplemental Fig. 2C). Mass spectrometry analysis verified that the major cobamide extracted from cultures grown with ALA is pseudocobalamin (Supplemental Fig. 2A, B).

**Figure 5.**
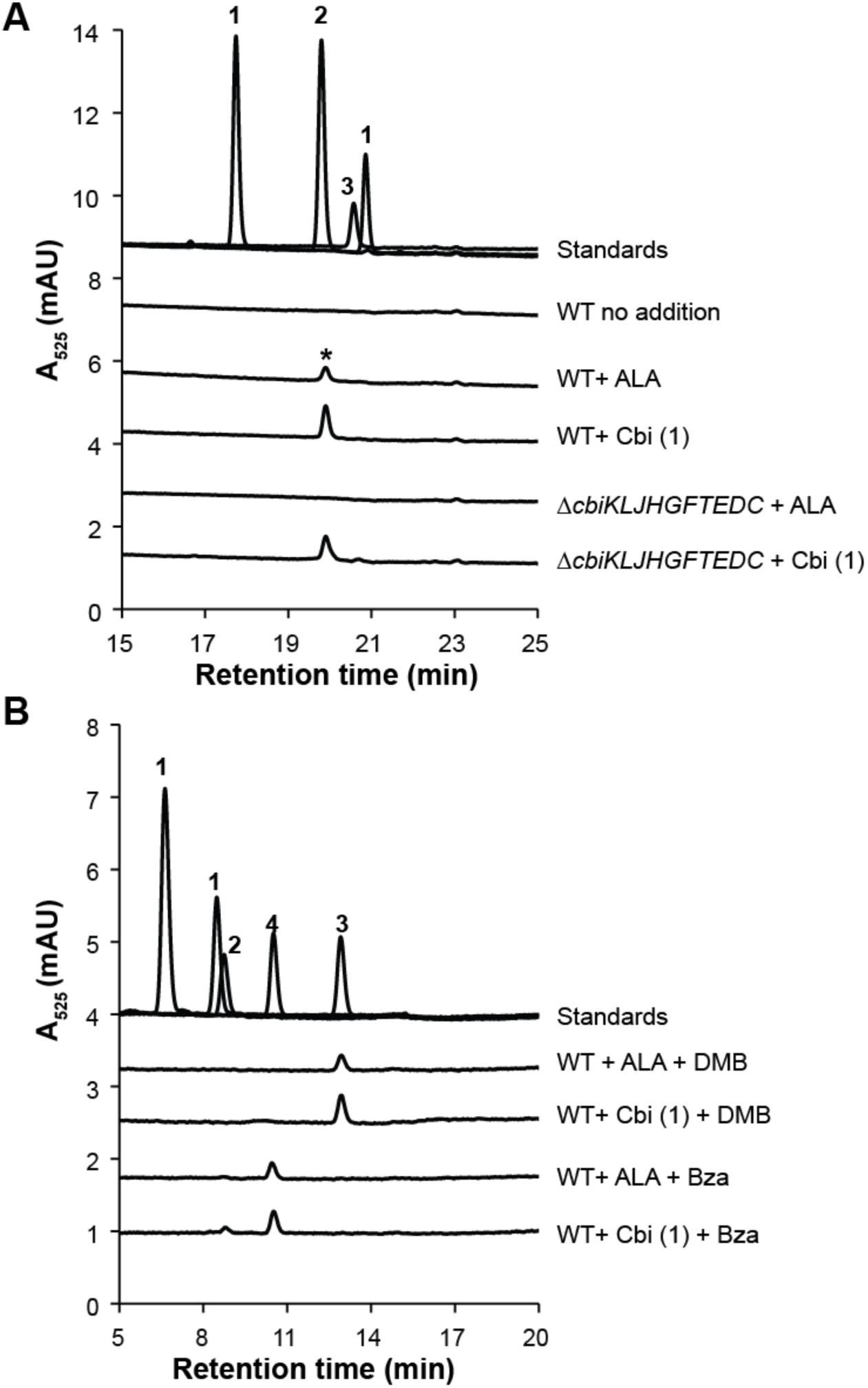
HPLC analysis of corrinoid extracts from *C. difficile* cultures. **A.** HPLC analysis of corrinoid extracts from cell pellets of *C. difficile* 630 Δ*erm* (WT) and 630 Δ*erm* Δ*pyrE* Δ*cbiKLJHGFTEDC* grown to saturation in CDDM with glucose with either 100 nM ALA or 10 nM dicyanocobinamide (Cbi) added. An asterisk (*) indicates the corrinoid peak validated by mass spectrometry (Supplemental Fig. 2). **B**. HPLC analysis of corrinoid extracts of *C. difficile* 630 Δ*erm* grown with either 100 nM ALA or 10 nM Cbi and 100 nM lower ligand bases DMB or Bza. An Agilent Eclipse Plus C18 column and an Agilent Zorbax SB-Aq column were used to separate corrinoid extractions in panels A and B, respectively. Cbi (1), pseudocobalamin (2), cobalamin (3), and [Bza]Cba (4) are shown as standards.

### *C. difficile* can perform guided biosynthesis but does not remodel cobamides

Some bacteria can perform guided biosynthesis, a process in which an exogenously provided, non-native lower ligand base is incorporated into a cobamide (32, 36, 48, 60, 61). To test if *C. difficile* is capable of guided biosynthesis to produce cobamides other than its native pseudocobalamin, either DMB (the lower ligand of cobalamin, Fig. 1A) or a related compound, benzimidazole (Bza, Fig. 1B) was added to cultures containing either ALA or Cbi. Analysis of corrinoid extracts showed that *C. difficile* could attach either of these exogenous lower ligands to form cobalamin and [Bza]Cba, respectively, with both precursors (Fig. 5B). A small amount of pseudocobalamin was also recovered in cultures containing Cbi with Bza (Fig. 5B).

Some bacteria and archaea are able to remodel cobamides by removing the lower ligand and nucleotide loop with the amidohydrolase enzyme CbiZ and rebuilding the cobamide with a different lower ligand (31, 62–64). We were unable to identify a *cbiZ* homolog in the *C. difficile* genome, and accordingly, we did not observe evidence of remodeling; when cobalamin, [2-MeAde]Cba, or [Cre]Cba was provided to *C. difficile*, the same cobamides were recovered from the cells (Fig. 6A).

**Figure 6.**
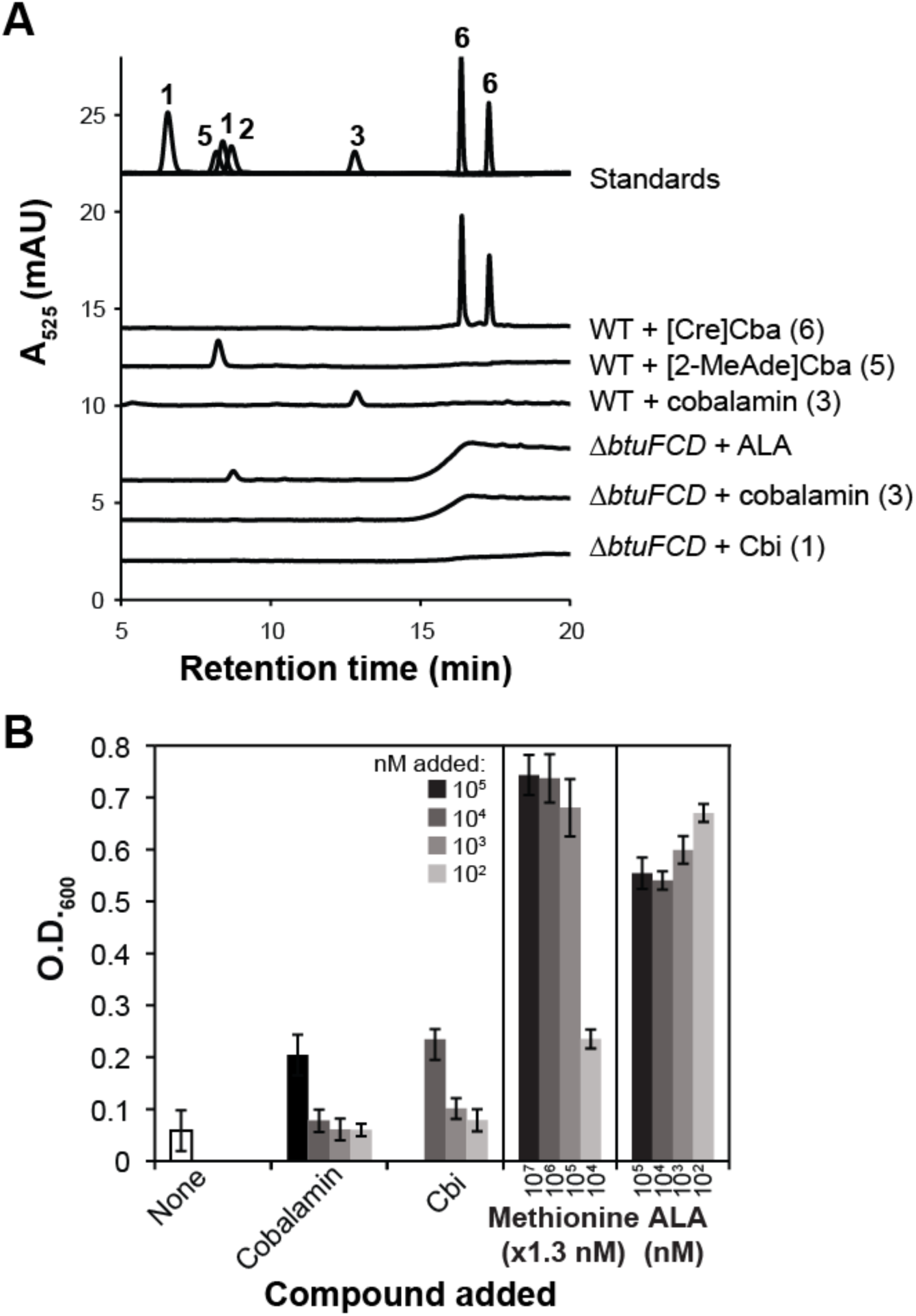
*C. difficile* Δ*btuFCD* mutant is impaired in cobamide and Cbi uptake. **A.** HPLC analysis of corrinoid extracts from cell pellets of *C. difficile* 630 Δ*erm* (WT) and *C. difficile* 630 Δ*erm* Δ*pyrE* Δ*btuFCD* grown with 10 nM cobamides or 100 nM ALA. Cbi (1), pseudocobalamin (2), cobalamin (3), [2-MeAde]Cba (5), [Cre]Cba (6) are shown as standards. **B.** Growth of *C. difficile* 630 Δ*erm* Δ*pyrE* Δ*btuFCD* in MetH-dependent conditions. The O.D._600_ of saturated cultures (23.5 hours) in CDDMK without methionine plus glucose and uracil is plotted as a function of the amount of compound added. Bars and error bars are the mean and standard deviation of three biological replicates.

### *C. difficile* requires *btuFCD* for efficient uptake of cobamides and Cbi

The presence of cobamides in the cellular fraction of cultures grown with either Cbi or a cobamide at nanomolar concentrations (Fig. 5, 6A) suggested that *C. difficile* takes up Cbi and cobamides via an active transporter. We identified a candidate cobalamin uptake operon (*btuFCD*) downstream of a sequence annotated as a cobalamin riboswitch, suggesting that these genes function in corrinoid import (27, 28, 65–70). No corrinoids could be detected in the cellular fraction of the *ΔbtuFCD* mutant grown with 10 nM Cbi or cobalamin (Fig. 6A). In contrast, ALA uptake is apparently unaffected in the *ΔbtuFCD* mutant, as pseudocobalamin can be recovered from the cellular fraction when ALA is provided (Fig. 6A). Furthermore, the Δ*btuFCD* mutant grew poorly in methionine-free medium even when Cbi or cobalamin was added at concentrations 10^3^ to 10^4^-fold higher than required for growth of the parental strain (Fig. 6B). The ability of methionine or ALA to support growth remained unaffected by the Δ*btuFCD* mutation (Fig. 6B). Interestingly, genomic analysis identified strains of *C. difficile* that contain a *tlpB* transposon insertion in *btuC*, likely rendering the BtuFCD transporter nonfunctional (Supplemental Fig. 3) (71). Of the genomes analyzed, the *tlpB* insertion in this locus appears to be restricted to strains in the PCR-ribotype 027 (RT027) clade, including the hypervirulent strain R20291, based on a multi-locus sequence typing (MLST) tree of *C. difficile* strains (Supplemental Fig. 3, red labels). This observation suggests that unlike strain 630 Δ*erm* examined in this study, members of the RT027 clade may be unable to take up cobamides and Cbi efficiently.

## Discussion

The potential of *C. difficile* to cause disease is closely linked to its ability to fill ecological niches made available by gut microbiota dysbiosis (13), using a suite of metabolic pathways to make use of newly available nutrient sources. *C. difficile* has an unusually high number of cobamide-dependent metabolisms encoded in its genome (25), but their functions have been underexplored. Here, we show that *C. difficile* is able to use many cobamides and cobamide precursors in two of its seven cobamide-dependent pathways. The promiscuous use of cobamides and the ability to bypass these cobamide-dependent metabolisms highlights the metabolic flexibility of *C. difficile.*

The cobalamin-dependent methionine synthase, MetH, is the most abundant cobamide-dependent enzyme in bacterial genomes (25) and is found in numerous organisms in all three domains of life, including humans (24). Compared to the majority of other MetH homologs that have been studied, our results indicate that the *C. difficile* MetH homolog is unusually promiscuous in its cobamide selectivity. For example, several eukaryotic algae grew robustly under MetH-dependent conditions with cobalamin, but did not grow with pseudocobalamin at the same concentrations (33). The human gut commensal bacterium *Bacteroides thetaiotaomicron* could use benzimidazolyl and purinyl cobamides for MetH-dependent growth, but could not use phenolyl cobamides (28). An example of MetH selectivity *in vitro* was in *Spirulina platensis*, where the purified enzyme bound its native cobamide, pseudocobalamin, with a higher affinity than cobalamin (72). An exception to this observed selectivity is another gut pathogen, *Salmonella enterica*, which can use its native cobamide, pseudocobalamin, in addition to cobalamin, [Phe]Cba, and [Cre]Cba for MetH-dependent growth, although other cobamides were not tested (48, 49). The versatility of *C. difficile*’s cobamide use is notable given the diversity of cobamides that have been detected in the gut (38).

In contrast to MetH, our growth experiments indicate that the selectivity of *C. difficile*’s NrdJ is more similar to that of other organisms that rely on NrdJ for growth. For example, *Sinorhizobium meliloti* was unable to grow with [Cre]Cba and grew poorly with pseudocobalamin relative to its native cobamide, cobalamin (36); *Lactobacillus leichmannii* could only use benzimidazolyl or purinyl cobamides (51); and *Euglena gracilis* grew well with cobalamin and [Bza]Cba and poorly with pseudocobalamin, [5-OHBza]Cba, and [Cre]Cba (33, 52). Unlike MetH, the NrdJ enzyme requires cobamides that can adopt the “base-on” configuration in which the lower ligand base is coordinated to the cobalt ion throughout the catalytic cycle (24). Phenolyl cobamides are unable to adopt the base-on configuration, so their inability to support growth in the NrdJ-dependent condition was expected. *C. difficile* 630 Δ*erm* also contains an active class III cobamide-independent RNR, NrdDG, which may be an important strategy to maintain deoxyribonucleotide synthesis when cobamides are scarce. However, in other species, under certain conditions the class II RNR provides an advantage over other RNR classes, such as during oxidative stress (73), although the conditions where NrdJ would provide an advantage for *C. difficile* have yet to be uncovered.

Seven different cobamides and the precursor Cbi have been detected in human feces (38). In stool samples of individuals not taking cobalamin supplements, the average total corrinoid present is approximately 1300 ng per gram feces, roughly equivalent to 1 µM (38). Cbi is found at tens of ng per gram (38). Growth experiments under MetH and NrdJ-dependent conditions showed that *C. difficile* 630 Δ*erm* reached maximum growth yield with as little as 1 nM cobamide or Cbi (Fig 3, 4). Based on the absence of corrinoids in the cellular fraction of a 630 Δ*erm* Δ*pyrE* Δ*btuFCD* strain (Fig. 6), we infer that strains with an insertion in *btuC* (Supplemental Fig. 3), including the hypervirulent R20291 and CD196 strains (71), would require cobamides or Cbi at extracellular concentrations higher than 100 µM if relying on cobamide-dependent enzymes. This suggests that these strains may not be able to use the cobamides or Cbi present in the gut.

Our results show that not only is *C. difficile* able to use multiple cobamides to support its metabolism, but it can also use the early precursor ALA to produce pseudocobalamin. The ability to use ALA to produce a cobamide, and thus not strictly rely on cobamide or Cbi uptake, could be important to strains with a transposon insertion in the *btuC* gene (70, 74). ALA concentrations in the human gut have not been reported. However, we speculate that, similar to cobamides and Cbi, ALA and possibly other early cobamide precursors could be provided by other members of the microbiota. Alternatively, ALA could be provided by the host either through the diet or via biosynthesis of heme, which also uses ALA as a precursor. Other commensal gut microbiota have been reported to be able to salvage ALA (25), suggesting that ALA could be available in the gut.

*C. difficile* is also able to incorporate non-native lower ligands to form benzimidazolyl cobamides (guided biosynthesis). Free benzimidazole bases have been found in animal gastrointestinal tracts such as rumen fluid and termite guts (75), but benzimidazole levels in the human gut have not been measured. The cobamides used by *C. difficile* could therefore also vary with the presence of different benzimidazole-producing organisms in the microbiota. Our results show that pseudocobalamin and most benzimidazolyl cobamides support growth of *C. difficile* equally for the two metabolisms we investigated in this study, but the cobamide preferences of the other five cobamide-dependent metabolisms have not been investigated.

We have identified cobamides and precursors that *C. difficile* can use *in vitro*, but which cobamides or cobamide precursors it predominantly uses in the gut remains to be discovered. Evidence from transcriptomics is ambiguous with respect to expression of genes encoding cobamide-dependent enzymes or cobamide biosynthesis during infection, likely due to differences in study design (15, 43, 76–78). Since both diet and the microbiota can contribute to the cobamide profile in the gut (38, 79, 80), the availability of cobamides may vary significantly across infection systems and affect the expression and use of cobamide biosynthesis and cobamide-dependent pathways by *C. difficile*. In one study, *hemB*, which encodes the enzyme that converts ALA to the next intermediate, porphobilinogen, was among the most highly expressed genes in *C. difficile* strain VPI 104363 in a mouse model (43), suggesting that *C. difficile* produces cobamides from ALA in the gut. How the cobamide content in the gut environment changes during *C. difficile* infection is unknown, but since much of the cobamide content in the lower gastrointestinal tract is produced by resident gut microbes (79, 80), it is possible that cobamide abundances change during dysbiosis. Further *in vivo* studies are needed to determine the extent to which cobamide metabolism is important to *C. difficile* associated disease.

## Materials and Methods

### Bacterial strains and growth conditions

*C. difficile* 630 Δ*erm*, an erythromycin-sensitive derivative of the isolate 630 (81), and *C. difficile* 630 Δ*erm* Δ*pyrE*, a derivative of 630 Δ*erm* with a uracil auxotrophy (50), were streaked from frozen stocks onto BHIS agar (82) before being transferred to *Clostridium difficile* defined medium (CDDM) containing casamino acids (83) and 8 g/L glucose. Agar plates and 96-well plates containing liquid cultures were incubated at 37°C in an anaerobic chamber (Coy Labs) containing 10% H_2_, 10% CO_2_, and 80% N_2_. For *C. difficile* 630 Δ*erm* Δ*pyrE* and derived strains, 5 µg/ml uracil was included in all defined media. For corrinoid extractions and NrdJ phenotype experiments, strains were cultured in CDDM plus 8 g/L glucose. For MetH phenotype experiments, CDDMK medium plus 8 g/L glucose without methionine was used. CDDMK contains the same salts, trace metals, and vitamins as CDDM, but the casamino acids, tryptophan and cysteine are replaced with the individual amino acids as follows: 100 mg/L histidine, 100 mg/L tryptophan, 100 mg/L glycine, 100 mg/L tyrosine, 200 mg/L arginine, 200 mg/L phenylalanine, 200 mg/L threonine, 200 mg/L alanine, 300 mg/L lysine, 300 mg/L serine, 300 mg/L valine, 300 mg/L isoleucine, 300 mg/L aspartic acid, 400 mg/L leucine, 500 mg/L cysteine, 600 mg/L proline, 900 mg/L glutamic acid (40). All liquid defined media were prepared by boiling under 80% N_2_/20% CO_2_ gas. After the pH stabilized between 6.8 and 7.2, the medium was dispensed into stoppered tubes and autoclaved. Filter-sterilized glucose and vitamins were added after autoclaving. Cultures in stoppered tubes were incubated at 37°C.

For MetH phenotype assays, *C. difficile* 630 Δ*erm* was grown in CDDM, then washed twice in CDDMK without methionine prior to inoculation in CDDMK at an optical density (O.D._600_) of 0.01 in a 96-well plate. For NrdJ phenotype assays, *C. difficile* 630 Δ*erm* Δ*pyrE* Δ*nrdDG* was grown in CDDM with 5 µg/ml uracil and 10 nM cyanocobalamin, and washed three times in CDDM medium without cyanocobalamin prior to inoculation in CDDM at an O.D._600_ of 0.01 in a 96 well plate. O.D._600_ was measured on a BioTek Synergy 2 plate reader after 23 to 24 hours of growth.

ALA, Cbi and cyanocobalamin were purchased from Sigma Aldrich. Other cobamides were purified from bacterial cultures as described in Men *et al*. (84)

### Strain and plasmid construction

The allelic coupled exchange (ACE) system of Ng *et al*. was used for construction of *C. difficile* mutant strains (50). Briefly, 500-1000 bp sequences flanking the target gene(s) (arms of homology) in the *C. difficile* 630 Δ*erm* genome (CP016318, https://www.ncbi.nlm.nih.gov/nuccore/CP016318.1/) were amplified by PCR (Supplemental Table 1) and then were cloned into pMTL-YN3 (Chain Biotech) by Gibson assembly (85) in *E. coli* XL1-Blue. Plasmid inserts were sequenced by Sanger sequencing before transformation of the plasmid into *E. coli* CA434 (Chain Biotech). Conjugation of *E. coli* CA434 and *C. difficile* 630 Δ*erm* Δ*pyrE* was performed as described (86), except that *C. difficile* and *E. coli* were each cultured for 5-8 hours prior to pelleting *E. coli* and mixing with the *C. difficile* recipient. After 16 hours growth on BHIS agar, the mixed cells were resuspended in 1 ml PBS, and 100 µl of the suspension was plated on each of 5-7 plates of BHIS agar with added 10 µg/ml thiamphenicol, 250 µg/ml D-cycloserine, and 16 µg/ml cefoxitin. Colonies were purified at least twice by streaking onto BHIS with 15 µg/ml thiamphenicol, 250 µg/ml D-cycloserine, and 16 µg/ml cefoxitin, before counterselection on CDDM agar supplemented with 2 mg/L 5-fluoroorotic acid (5-FOA) and 5 µg/ml uracil. The resulting colonies were purified by streaking at least twice on the counterselection medium prior to screening by colony PCR for the deletion and the presence of the *C. difficile* toxin gene *tcdB* (86). For the deletion of *nrdDG*, 10 nM cobalamin was added to all media during the ACE procedure.

### Corrinoid extraction and analysis

*C. difficile* was grown in 50 ml CDDM plus 8 g/L glucose under 80% N_2_/20% CO_2_ headspace for 16-22 hours at 37°C prior to corrinoid extraction. Two cultures were combined for each condition for a total volume of 100 ml for each extraction. Corrinoid extractions were performed as described (31), except that cell pellets were autoclaved for 35 minutes and cooled prior to addition of methanol and potassium cyanide. Two or more biological replicates were performed for each condition.

High-performance liquid chromatography (HPLC) analysis was performed with an Agilent Series 1200 system (Agilent Technologies, Santa Clara, CA) equipped with a diode array detector with detection wavelengths set at 360 and 525 nm. For Fig. 5B and 6A samples were injected onto an Agilent Zorbax SB-Aq column (5 µm, 4.6 × 150 mm) at 30°C, with 1 mL/min flow rate. Compounds in the samples were separated with a gradient of 25 to 34% acidified methanol in acidified water (containing 0.1% formic acid) over 11 min, followed by a 34 to 50% gradient over 2 min, and 50 to 75% over 9 min. For Fig. 5A, samples were injected onto an Agilent Eclipse Plus C18 column (5 µm, 9.6 × 250 mm) at 30 °C, with 2 mL/min flow rate. Compounds in the samples were separated with a gradient of 10 to 42% acidified methanol in acidified water over 20 min. The amount of standards that were injected were as follows: Cbi (**1**), 200 pmol; pseudocobalamin (**2**), 225 pmol; cobalamin (**3**), 50 pmol; [Bza]Cba (**4**), 114 pmol; [2-MeAde]Cba (**5**), 114 pmol; [Cre]Cba (**6**), 151 pmol. 5% to 20% by volume *C. difficile* samples were injected.

**Table 1:**
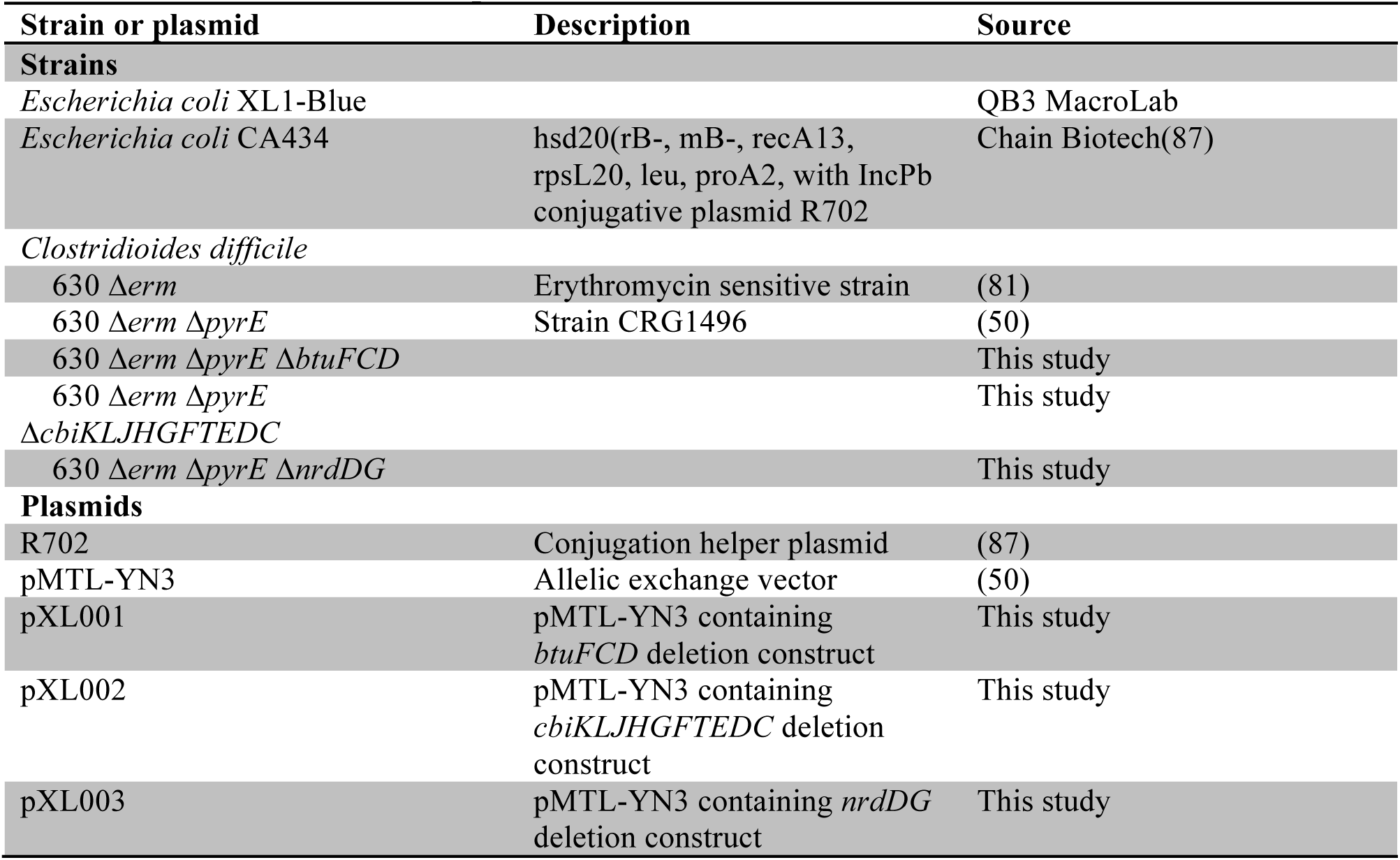
Bacterial strains and plasmids.

## Supplemental material

Supplemental methods, figures, and tables are provided.

## Acknowledgements

This work was supported by grants NIH DP2AI117984 and R01GM114535 to M.E.T.

We thank Aimee Shen for her extensive advice and protocols for working with *C. difficile*, Craig Ellermeier for advice on working with *C. difficile*, and John Taylor, Gary Andersen, Christian Sieber, and Eric Dubinsky for advice on creating the phylogenetic tree. *C. difficile* 630 Δ*erm* and 630 Δ*erm* Δ*pyrE* were provided to us by Andrew Hryckowian with permission from Nigel Minton. We thank Olga Sokolovkaya and Kenny Mok for help with HPLC analysis and Dr. Rita Nichiporuk at the QB3/Chemistry Mass Spectrometry Facility at UC Berkeley for performing the mass specrometry analysis. We thank Luis Valentin-Alvarado for helpful discussions about *C. difficle* metabolism and genetic tools. We also thank Ellen Simms, Olga Sokolovskaya, Sebastian Gude, Gordon Pherribo, and Kenny Mok for critical reading of this manuscript.

## Author Contributions

A.N.S. performed growth experiments, corrinoid extractions, and phylogenetic analysis. X.L. and A.N.S. created the mutant strains. A.N.S. and M.E.T. wrote the manuscript.

