## Supplemental materials for "Flexible cobamide metabolism in *Clostridioides (Clostridium) difficile* 630 Δ*erm*"

### Supplemental Methods

#### Cobamide purification and mass spectrometry analysis

The peak co-eluting with pseudocobalamin of one corrinoid extraction of 100 ml of *C. difficile* 630 Δ*erm* grown for 23.5 hours in CDDM with 10 µM ALA was purified using the Agilent Eclipse Plus C-18 column (5 µm, 9.6 × 250 mm) at 30 ºC, with 2 mL/min flow rate. Compounds in the samples were separated with a gradient of 10 to 42% acidified methanol in acidified water (containing 0.1% formic acid) over 20 min. The collected peak was desalted on a C-18 SepPak column (Waters), dried, and resuspended in 100 µl water. The UV-Vis spectra of the sample and a pseudocobalamin standard were collected during the HPLC separation. This sample was additionally purified on an Agilent Zorbax SB-Aq column (5 µm, 4.6 × 150 mm) at 30°C, with 1 mL/min flow rate. Compounds in the samples were separated with a gradient of 25 to 34% acidified methanol in acidified water (containing 0.1% formic acid) over 11 min, followed by a 34 to 50% gradient over 2 min, and 50 to 75% over 9 min. The collected peak was desalted and dried. The dried sample was resuspended in MeOH to 20-40 uM, and analyzed on a Finnigan LTQ FT mass spectrometer (Thermo Fischer Scientific) equipped with electrospray ionization (ESI) source in positive ion mode at the UC Berkeley QB3/Chemistry Berkeley Mass Spectrometry facility. Xcalibur™ software (version 2.0.7, Thermo) was used for both data acquisition and data analysis.

#### *C. difficile* MLST tree construction

For the 248 *C. difficile* genomes classified as "finished" or "permanent draft" in the JGI/IMG database (1) (accessed March 2019, https://img.jgi.doe.gov/cgi-bin/mer/main.cgi), the seven MLST gene sequences, *adk, atpA, dxr, glyA, recA, sodA,* and *tpi* (2), were downloaded and aligned individually using MUSCLE (3). The alignments were concatenated and genomes missing one or more MLST genes or having duplicate genes were removed from the analysis, resulting in a total of 79 strains analyzed. The concatenated alignment was manually trimmed in UGENE (4), and columns with 95% or greater gaps were removed with trimAL (5). This alignment was used as input for RAxML 8.2.12 (6) on the CIPRES webserver (https://www.phylo.org/) (7) with 100 bootstraps, using the GTRCAT model. The tree was visualized and annotated in iTOL (https://itol.embl.de/) (8).

### Supplemental Figures and Tables

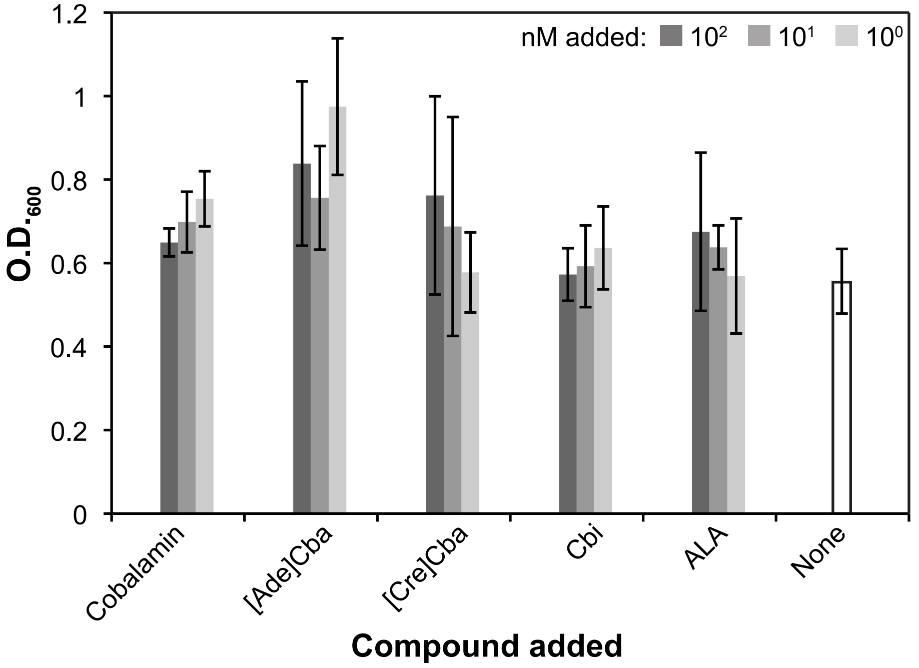


**Supplemental Figure 1: Growth yield of *C. difficile* 630 Δ*erm* Δ*pyrE* does not change in response to external cobamides when methionine is present**. *C. difficile* 630 Δ*erm* Δ*pyrE* was grown in CDDM casamino acid medium plus uracil and glucose for 23.5 hours. The mean $OD_{600}$ of three biological replicates is plotted. Error bars are the standard deviation.

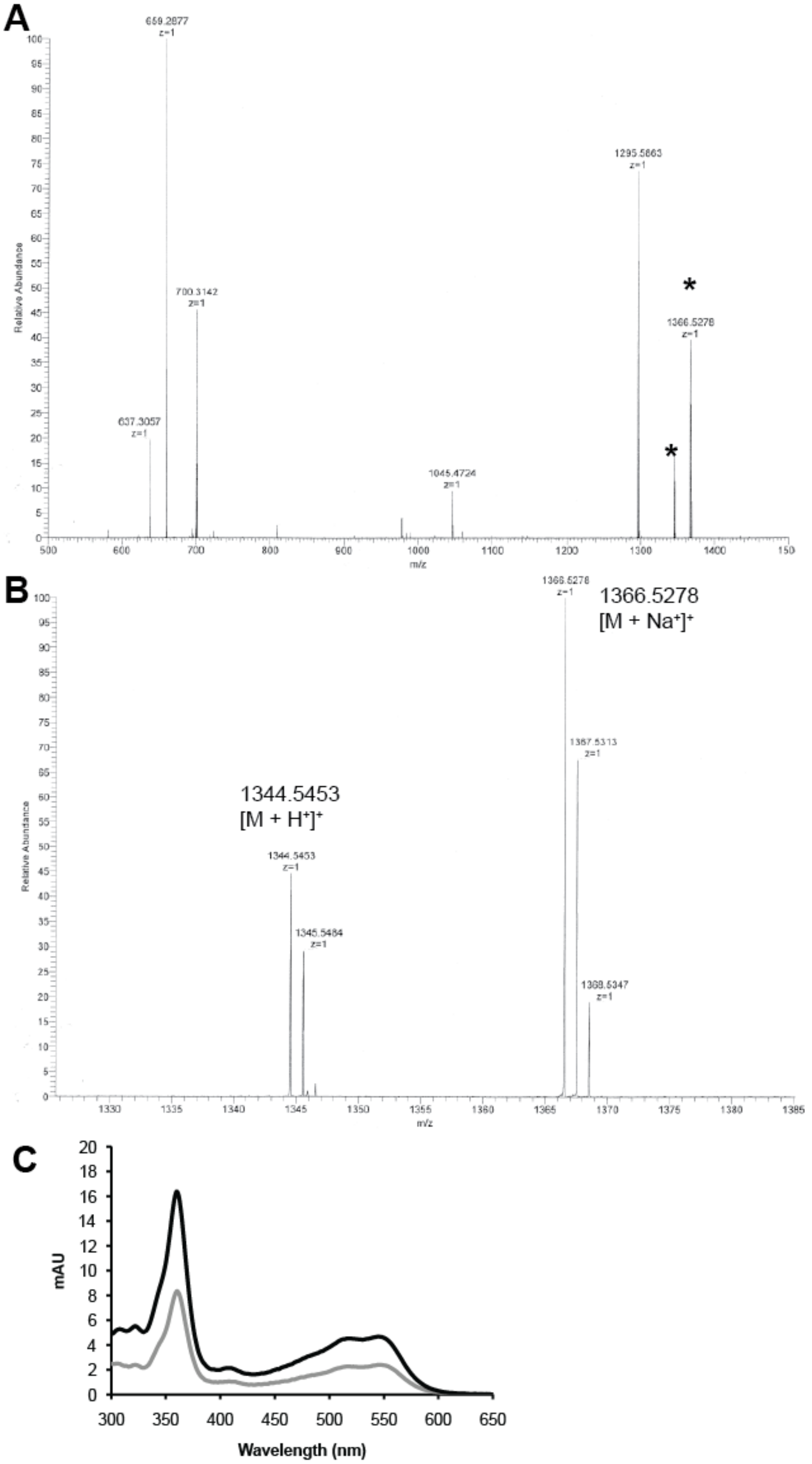


**Supplemental Figure 2.** Characterization of the cobamide produced by *C. difficile* with exogenous ALA. **A.** ESI-MS trace of range scanned (500-1800), with asterisks indicating cobamide peaks. Other peaks present were not identified. **B.** Subset of the scan to show the cobamide peaks. The expected molecular formula for cyanopseudocobalamin, $C_{59}H_{83}Co_1N_{17}O_{14}P_1$, was confirmed. **C.** UV-Vis spectrum of the material purified by HPLC for mass spectrometry analysis (black) and a cyanopseudocobalamin standard (gray).

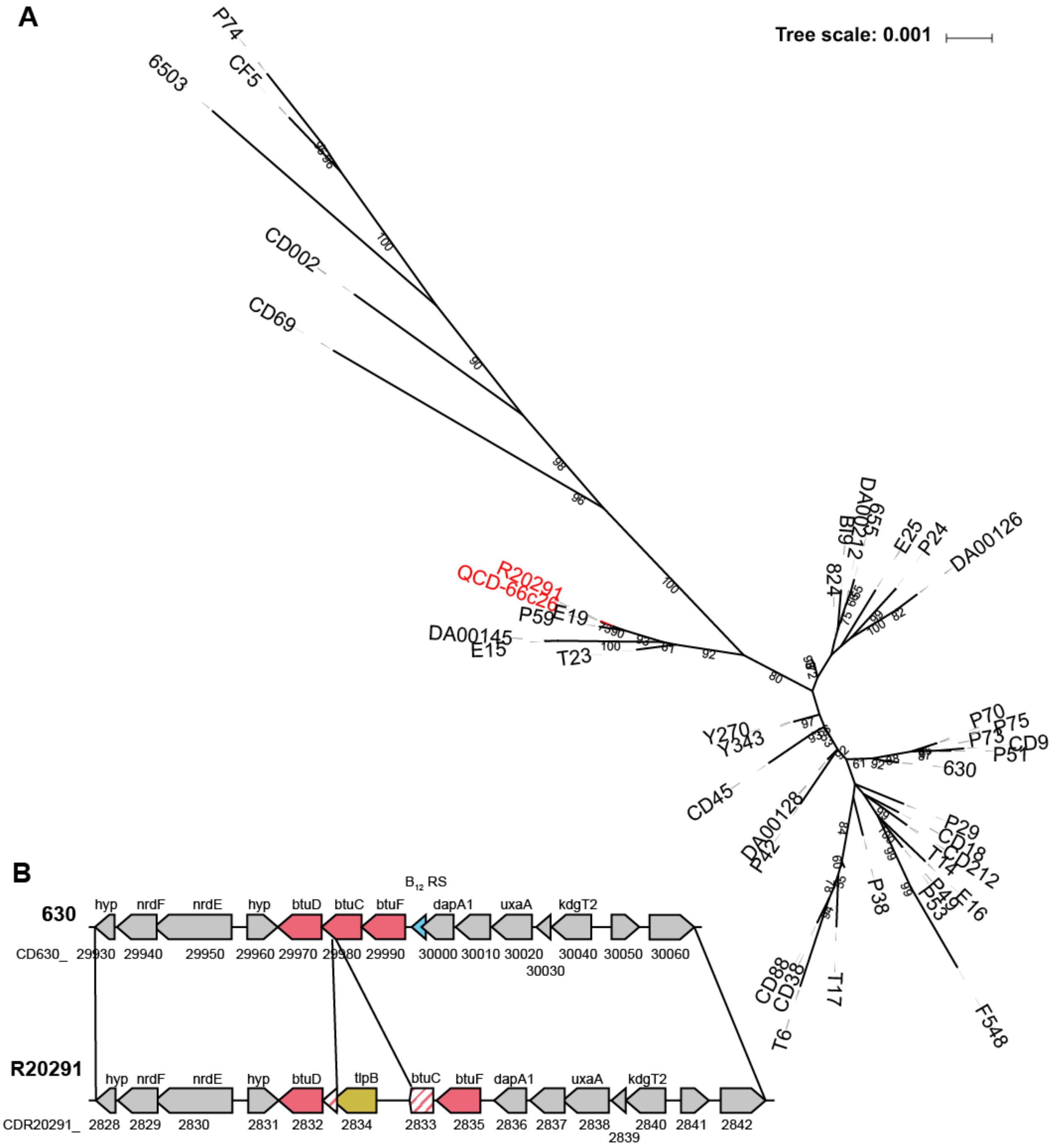


**Supplemental Figure 3. Distribution of *tlpB* transposon insertion in *btuC* in C. *difficile* strains A.** Maximum likelihood multi-locus sequence typing (MLST) tree of 79 *C. difficile* strains labeled with their strain designation. Branch labels are support values from 100 bootstraps. Genomes with the *tlpB* insertion in *btuC* are labeled in red, those without the *tlpB* insertion in *btuFCD* are in black. Clades with average branch lengths less than 0.0001 substitutions per site have been collapsed and bootstrap values less than 50 have been removed to improve readability. **B.** Gene neighborhood diagram of strain 630, which lacks the *tlpB* insertion, and strain R20291, which has a *tlpB* insertion in *btuC* (shown in yellow). The *btuC* pseudogene is indicated by pink stripes. Gene names and locus tags are shown.

**Supplemental Table 1: Primers used in this study**

| Primer ID | Sequence | Purpose |
|---|---|---|
| P2188 | CAT AAT ATG TCA GAG AAT ACT GTA GTC | *tcdB* presence |
| P2189 | GTT CTG AGG TAT ATT CTG GTA TAT ATT | *tcdB* presence |
| P2269 | GCT TGA TGT GTT GGT AGC AC | Checking pMTL-YN3 for insert |
| P2270 | AAG TAC ATC ACC GAC GAG CA | Checking pMTL-YN3 for insert |
| P2271 | AAC AGC TAT GAC CGC GGC CGC ATT TTA ATG AAA ACT ATT TC | Gibson assembly primer for pMTL-YN3 and *btuFCD* arms of homology |
| P2272 | TTC AAA AAA ATT ATA ATC TAT ACT CTA ATT TAT TGT TGA CCT CTT TGC AAG G | Gibson assembly primer for pMTL-YN3 and *btuFCD* arms of homology |
| P2273 | CCT TGC AAA GAG GTC AAC AAT AAA TTA GAG TAT AGA TTA TAA TTT TTT TGA A | Gibson assembly primer for pMTL-YN3 and *btuFCD* arms of homology |
| P2274 | CAG GCC TCG AGA TCT CCA TGG TAT GGA TAT GCA AAA AGA AC | Gibson assembly primer for pMTL-YN3 and *btuFCD* arms of homology |
| P2282 | ACA GCT ATG ACC GCG GCC GCC TAA TTT CTA TAG CTA AAG C | Gibson assembly primer for pMTL-YN3 and c*biKLJHGFTEDC* arms of homology |
| P2286 | AGT CTC CTT TAA ATA TTG CTT TCA CTT ATG TAT TCC ATT CTA TTT CCC CCT TAA T | Gibson assembly primer for pMTL-YN3 and c*biKLJHGFTEDC* arms of homology |
| P2287 | ATT AAG GGG GAA ATA GAA TGG AAT ACA TAA GTG AAA GCA ATA TTT AAA GGA GAC T | Gibson assembly primer for pMTL-YN3 and *cbiKLJHGFTEDC* arms of homology |
| P2288 | AGG CCT CGA GAT CTC CAT GGA TAC CTG TTG GAA AAG GAA T | Gibson assembly primer for pMTL-YN3 and *cbiKLJHGFTEDC* arms of homology |
| P2289 | ACA GCT ATG ACC GCG GCC GCC AAT AAG TTT TTT ACA GAT T | Gibson assembly primer for pMTL-YN3 and *nrdDG* arms of homology |
| P2290 | CAA TAT ATA GTA ACA GGA GGT TTT TTT AAA ATA TAA ATA AAC AGG ATT AAA TAT ATG C | Gibson assembly primer for pMTL-YN3 and *nrdDG* arms of homology |
| P2291 | GCA TAT ATT TAA TCC TGT TAA TTT ATA TTT TAA AAA AAC CTC CTG TTA CTA TAT ATT G | Gibson assembly primer for pMTL-YN3 and *nrdDG* arms of homology |
| P2292 | TCT GCA GGC CTC GAG ATC TCC ATG GTA TTA CTA TAC CAA CTT TTT CTT TTA GAG T | Gibson assembly primer for pMTL-YN3 and *nrdDG* arms of homology |
| P2404 | GAA GGT GAT TTT AAT GAA AAC TAT TTC TAT TTC TAA ACA AG | Flanking primer for checking *btuFCD* knockout |
| P2405 | TAT GCA AAA AGA ACT TAT AGA TTT AGT AAC TAG TC | Flanking primer for checking *btuFCD* knockout |
| P2406 | CGT CCG TCT TAT CTA CTG ATT GAT AAT AAT C | Internal primer for checking *btuFCD* knockout |
| P2663 | GCA AAA TAT GAT TAC TTG ATG CCT TG | Flanking primer for checking *cbiKLJHGFTEDC* knockout |
| P2664 | TCT TAC AAG CAA CAC TGA AAT TAT G | Flanking primer for checking *cbiKLJHGFTEDC* knockout |
| P2451 | CAG GAT AAC TAA CCC AAT AAG GCT TTG GTA | Flanking primer for checking *nrdDG* |

| | | |
|---|---|---|
| | ATA AGA CTT C | knockout |
| P2452 | TGC TTT ATT TGT TTG CCC TTT TCT TGA GGA C | Flanking primer for checking *nrdDG* knockout |
| P2458 | GAA GAT ATA CAA GAT TCT GTA GTT AAG GTT C | Internal primer for checking *nrdDG* knockout |
| P2459 | AGA AGT ATC TGT TCC GAA GTT TAC ACT TG | Internal primer for checking *nrdDG* knockout |
